# Metabolites associated with type 2 diabetes and Alzheimer’s disease trigger differential intracellular signaling responses in mouse primary neurons

**DOI:** 10.1101/2025.04.23.650328

**Authors:** Brendan K. Ball, Madison K. Kuhn, Rebecca M. Fleeman Bechtel, Elizabeth A. Proctor, Douglas K. Brubaker

## Abstract

Alzheimer’s disease (AD) is a progressive neurodegenerative disease that is accelerated by the pathological features of type 2 diabetes (T2D). Neuroinflammation is an extensively studied component shared by T2D and AD that remains poorly understood. In this work, we studied nine blood-brain barrier permeable metabolites associated with protective or harmful effects of AD and T2D in literature (aminoadipic acid, arachidonic acid, asparagine, D-sorbitol, fructose-6-phosphate, lauric acid, L-tryptophan, niacinamide, and retinol) and quantified intracellular signaling responses in primary cortical neuron monocultures. After stimulation of neuronal cultures with each metabolite, we quantified signaling analytes with a Luminex assay. Using univariate and multivariate analysis approaches, we identified potential intracellular signaling pathways linked to AD and T2D pathology. With partial least squares discriminant analysis, we identified the separation between the disease and protective-associated metabolites. We identified Akt and STAT5 up-regulation by AD- and T2D-associated metabolites, whereas c-Jun and MEK1 were up-regulated by disease-protective metabolites. Finally, we performed a canonical correlation analysis to link neuronal cytokine data we previously collected from these cultures to our new intracellular signaling data, to which we found intracellular proteins associated with detrimental and protective properties that correlated with IL-9 and MCP-1, respectively. Our experimental and computational approach identified potential associations between intracellular and cytokine signaling molecules in the context of AD and T2D pathology. Nevertheless, primary neuron responses to metabolites associated with T2D and AD may contribute to neuroinflammation and progressive cognitive decline.

## INTRODUCTION

Alzheimer’s disease (AD) is a complex neurodegenerative disease that results in continuing memory loss and cognitive deficits. A major contributor to the development of AD is type 2 diabetes (T2D), which is characterized by persistent hyperglycemia and dysregulated insulin signaling. The onset of T2D, which often precedes decades before cognitive deficits^1^, enhances the risk of AD by more than 60% compared to individuals without T2D^2–4^. One possible explanation for the elevated risk of AD is the shared pathophysiological mechanisms between the two diseases^5^. A pathway shared between T2D and AD is immune dysregulation^6,7^. Our group^8–10^ and other researchers^11–13^ have identified shared chemokine signaling and relevant immune system pathways by which T2D may influence the development of AD.

A potential cause for the activation of neuroinflammatory pathways is through disruption to the blood-brain barrier (BBB), a highly protective membrane that selectively regulates the entry of beneficial nutrients and removal of biological waste^14,15^. The BBB is made up of a tightly bound formation of astrocytes, pericytes, and endothelial cells under healthy conditions, but these same cells degenerate and are lost under T2D^16^ and AD^17^ pathology, resulting in BBB leakage^18^. Prior studies have also reported a decrease in BBB integrity in rodent models fed with a high-fat diet^19^ and found cognitive dysfunction linked to diet-induced BBB disruption^20^.

The connection between T2D, AD, and BBB impairment was also reported in humans diagnosed with T2D and AD^21^. Therefore, the circulation of harmful molecules may induce further breakdown of the BBB and lead to downstream neuroinflammatory responses by local cells in the brain. This chronic, low-grade inflammation in the brain may contribute to cognitive decline and the development of AD.

A multitude of neuroinflammatory pathways are altered under conditions of T2D^22^ and AD^23^. Neuronal cells may release cytokine cues to be received by neighboring cells, or relay signals inside the cell to regulate and maintain biological function within their environment^24^. These signaling molecules are actively involved in triggering inflammatory responses^25^, modulating transcriptional changes^26^, and facilitating cellular communication^27^. Cytokines such as interleukins^28^ and chemokines^29^ mediate both pro- and anti-inflammatory responses and can modulate neuronal synaptic and glutamatergic function^30^. Intracellular pathways such as JNK^31^ (c-Jun N-terminal kinases) are major regulators of several cellular processes and can mediate neuroinflammation and contribute to the disruption of the BBB. Other pathways such as STAT1 (signal transducer and activator of transcription 1)^32^ have been associated with neuroinflammation and the eventual onset of AD. In a positive feedback-like loop BBB damage may allow additional harmful substances to enter the brain and cause further disruption to neuronal cells. Despite the overlap identified in high-level signaling pathways, the potential downstream mechanisms are broad and our understanding of how T2D affects AD progression in the context of neuroinflammation and cellular signaling is largely limited.

To understand this relationship between T2D and AD, we focused on the hypothesis that BBB-permeable metabolites associated with T2D or AD may induce neuroinflammatory changes in the brain and contribute to cognitive decline. To test this hypothesis, we stimulated murine primary neurons with T2D- and AD-associated metabolites and quantified signaling responses in a panel of intracellular signaling proteins to determine if neuronal cell responses differed across different metabolite groups that we identified to be associated with T2D or AD in literature. We integrated these data with our previously published cytokine data from the same cultured cells and experimental conditions^8^ via canonical correlation analysis (CCA). This newly generated and integrated data allowed us to characterize a relationship between intracellular signaling and cytokine secretion in connection to AD/T2D-associated metabolites.

## RESULTS

### Characterizing neuronal signaling responses to T2D- and AD-associated metabolites

We performed neuronal stimulation experiments using nine metabolites either associated with or protective against T2D or AD, including L-tryptophan, niacinamide, lauric acid, asparagine, retinol, aminoadipic acid, arachidonic acid, D-sorbitol, and fructose-6-phosphate (**Fig. 1a**).

**Figure 1.**
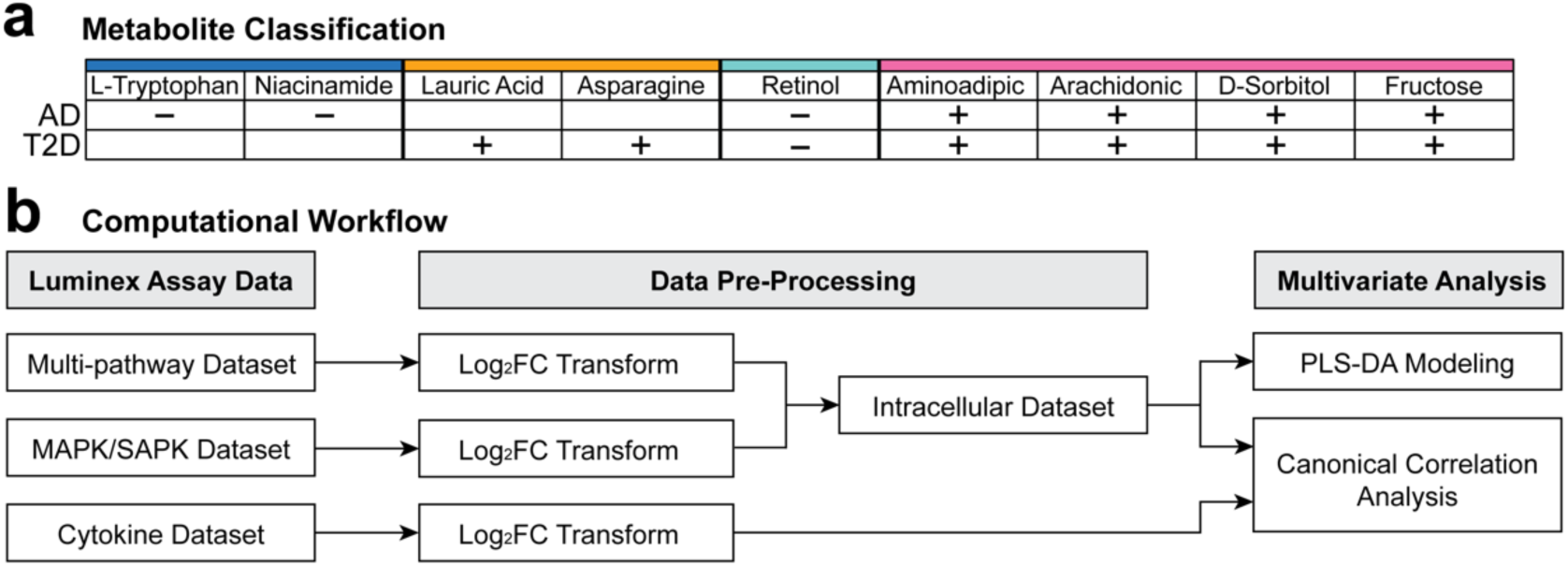
Metabolite identification and computational workflow. **(a)** Metabolites were identified through literature with associations to AD-protective, T2D, AD/T2D-protective, and both AD/T2D. A “+” is associated with disease, whereas a “–” is associated with protective conditions. Primary neurons were grown in 6-well plates and stimulated with metabolites and their respective vehicles and prepared for Luminex **(b)** After data acquisition, multi-pathway and MAPK/SAPK datasets were separately processed and combined into an intracellular signaling dataset. We performed multivariate analysis by creating partial least squares discriminant analysis models on the intracellular data, then correlated our cytokine and intracellular responses together with canonical correlation analysis.

Lauric acid^33,34^ and asparagine^35,36^ are metabolites positively associated with T2D (T2Dτ). For metabolites positively associated with both AD and T2D (AD/T2Dτ), we identified fructose-6-phosphate^37,38^, arachidonic acid^39,40^, aminoadipic acid^36,41^, and D-sorbitol^36,42^ with connections to freezing behavior, spatial memory impairment, and induction of glucose intolerance in rodent models. Metabolites with AD protective (ADτ) associations included niacinamide^43,44^ and L-tryptophan^45,46^, by which the presence of such metabolites was shown to reduce or prevent cognitive deficits. Retinol is a metabolite with dual T2D and AD protective (AD/T2Dτ) properties^33,47^, reported to be downregulated in humans in cases of AD and T2D.

We used an *in vitro* approach to collect and quantify neuronal intracellular responses to individual metabolites with T2D and/or AD associations. We generated primary cortical neuron monocultures from CD1 mouse embryos for metabolite stimulation. We stimulated neuron cultures with each metabolite (or its respective vehicle control) incubated for 72 h and quantified the levels of intracellular signaling proteins for downstream computational analysis (**Fig. 1b**).

### Neuronal JNK, MEK1, Akt, and STAT5 signaling differentially responded to T2D and AD associated metabolites

We merged two datasets (immunoassay kits MAPK/SAPK and multi-pathway) containing intracellular protein levels from the same biological replicates for a total of 16 different intracellular proteins for downstream computational analysis. The proteins included Akt, ATF2, c-Jun, CREB, ERK/MAPK ½, HSP27, JNK, MEK1, MSK1, NFkB, p38, p53, p70S6K, STAT1,

STAT3, and STAT5. We computed the z-score transformation onto the quantified total protein levels and hierarchically clustered by similarities across protein changes with respect to vehicle groups (**Fig. 2a**). Of the 16 intracellular proteins, we identified seven that contained significant response levels to the different metabolite conditions (Kruskal-Wallis test, *q* < 0.10). The significant proteins included STAT5 (*q* = 0.0651), Akt (*q* = 0.0950), JNK (*q* = 0.0950), STAT1 (*q* = 0.0950), p70S6K (*q* = 0.0762), MEK1 (*q* = 0.0651), and ATF2 (*q* = 0.0950).

**Figure 2.**
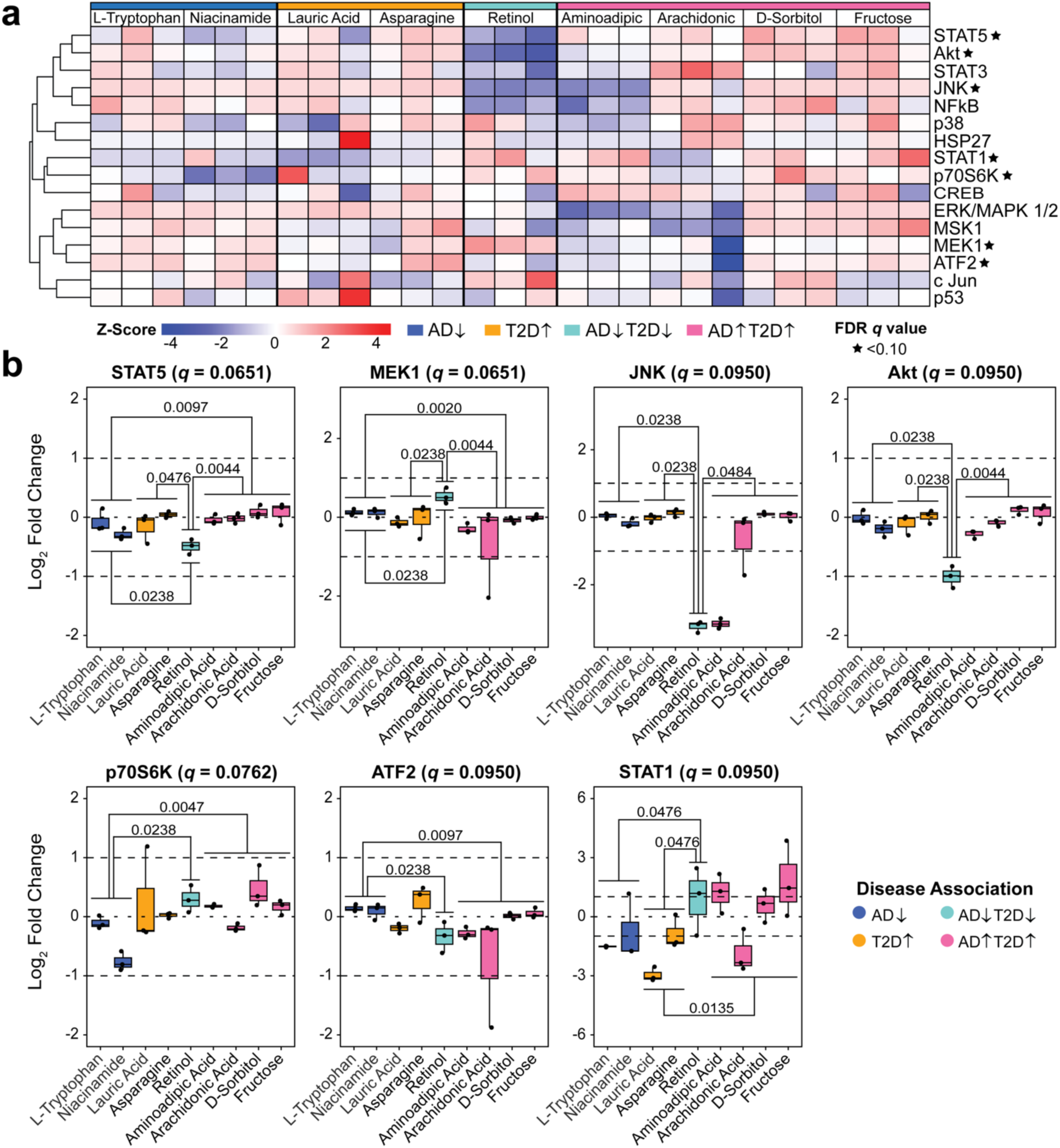
Hierarchical clustering of intracellular protein concentrations in response to metabolite treatments. **(a)** Intracellular proteins were hierarchically clustered by their respective z-scores. Significant pathway proteins (FDR *q* < 0.10) were denoted with a star as determined by a Kruskal-Wallis test adjusted by the Benjamini-Hochberg method. **(b)** Mann-Whitney pairwise testing of the log_2_ fold change (metabolite/vehicle) of significant proteins identified by the Kruskal-Wallis test. Significance between different metabolite condition pairs was established with a *p* < 0.05.

We performed a pairwise Mann-Whitney test on the seven intracellular proteins we deemed to be significant from the Kruskal-Wallis test (**Fig. 2b**) and identified JNK (c-Jun N-terminal kinase), MEK1 (mitogen-activated protein kinase 1), Akt (protein kinase B), and STAT5 (signal transducer and activator of transcription 5) to have significantly different responses between metabolites with proactive associations (ADτ and AD/T2Dτ) and the detrimental metabolites (T2Dτ and AD/T2Dτ), STAT5 and MEK1 had similar patterns of significant signaling responses concerning metabolite groups, except that the responses to retinol were opposite. Moreover, JNK and Akt demonstrated similar response patterns across all metabolite groups. The total protein abundance of p70S6K, ATF2, and STAT1, though significant by their *q* value, did not show a clear distinction between the protective and disease-associated metabolites. We also visualized the protein abundances and verified a lack of significance across the metabolite groups for proteins that were not considered significant by the Kruskal-Wallis test (**Supplementary Fig. S2**).

### MEK1 levels are linked to AD/T2D-protective, Akt, STAT3, and STAT5 to AD/T2D, JNK to T2D, and c-Jun to T2D-protective associations

We used PLS-DA, a multivariate dimensional reduction technique to identify protein biomarkers that may contribute to the separation of metabolite stimulation conditions to further understand the relationships among signaling responses and disease associations. We first constructed a four-way PLS-DA model, where metabolites were classified into AD-protective (ADτ), T2D-associated (T2Dτ), AD/T2D-protective (AD/T2Dτ), and both AD/T2D-associated (AD/T2Dτ).

We discriminated against the four metabolite conditions (ADτ, AD/T2Dτ, T2Dτ, and AD/T2Dτ) in the first two latent variables and found differential intracellular protein levels in response to the metabolites (**Fig. 3a**). We assessed whether proteins contributed more than average to the model’s classification by their variable importance in projection (VIP) score (VIP > 1) (**Fig. 3b**). For proteins contributing to ADτ, we found NFkB on LV1 and no proteins on LV2. For AD/T2Dτ, we found c-Jun on LV1, STAT1 on LV2, and MEK1 on both LVs. We identified JNK on both LV1 and LV2 and Erk/MAPK ½ on LV2 associated with T2Dτ. Finally, we identified Akt, STAT3, and STAT5 on both LV1 and LV2, which was associated with the final metabolite classification, AD/T2Dτ. Taken together, these findings show that intracellular proteins like MEK1, c-Jun, and STAT1 may drive protective associations (ADτ and AD/T2Dτ) whereas STAT3, STAT5, Akt, and JNK may drive disease-like conditions (T2Dτ and AD/T2Dτ).

**Figure 3.**
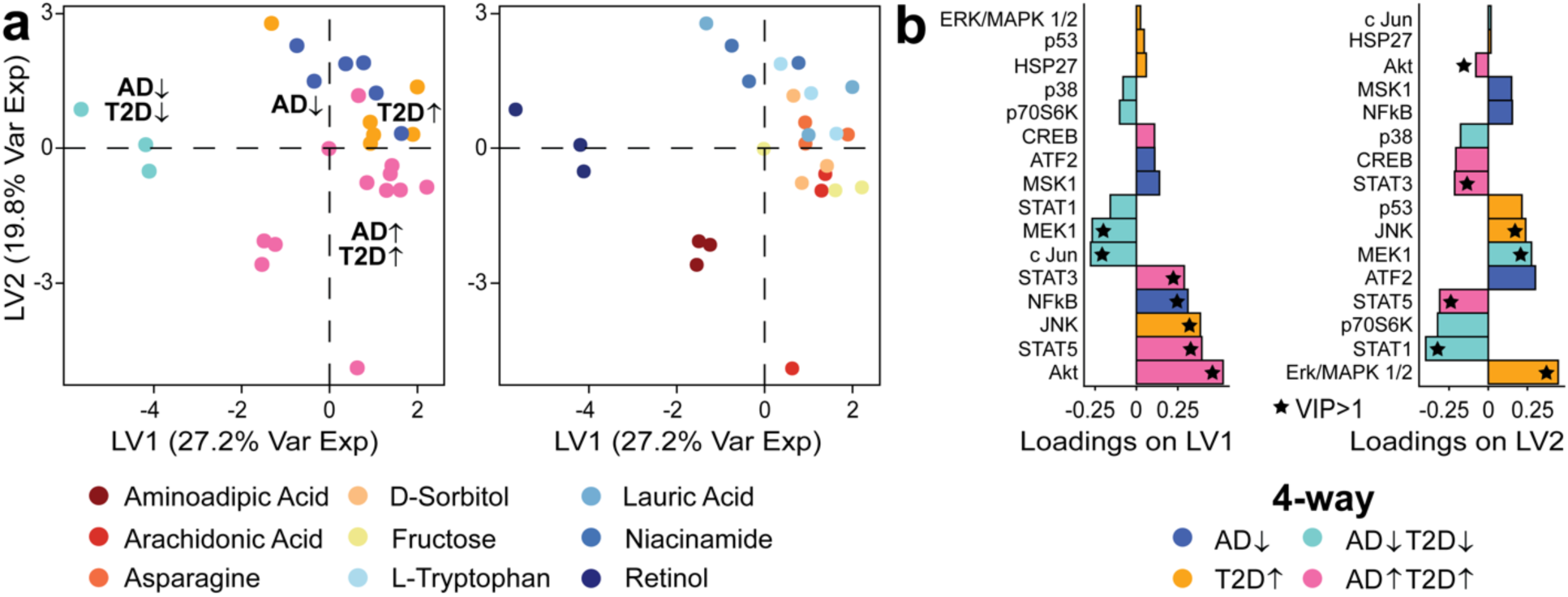
Intracellular protein levels are differentially abundant in response to metabolites associated with T2D and AD. **(a)** Four-way disease classification model annotated by disease status (left) and metabolite (right). **(b)** The loadings for latent variables 1 and 2 with a star indicate a VIP score greater than 1. The color marked on each variable in the loadings represents the analyte with the highest contribution to a metabolite condition.

Having identified differences in intracellular protein levels across our four-way PLS-DA model, we next determined if classifying our metabolites by AD/AD-protective or T2D/T2D-protective associations corroborated our findings. Our motivation stemmed from our recognition that these metabolites are complex molecules and may participate in multiple downstream biological processes, which may complicate how metabolite associations are defined. Thus, we classified metabolites by discriminating against AD (ADτ vs. ADτ) or T2D (T2Dτ vs. T2Dτ) association. In these models, metabolites that did not fit the AD- or T2D-only classification were excluded from such respective PLS-DA models.

In our model that associated metabolites as either AD-protective or AD, we found negative scores associated with AD and positive scores with AD-protective association (**Fig. 4a**). Metabolites associated with AD such as asparagine, fructose-6-phosphate, D-sorbitol, and aminoadipic acid triggered total protein levels of STAT5, Akt, STAT3, p70S6K, CREB, STAT1, HSP27, p38, JNK, and NFkB. Metabolites with an AD-protective association had higher levels of MEK1, Erk/MAPK ½, c-Jun, ATF2, p53, and MSK1. Encoded on AD LV1 that separated the two clusters, we found p70S6K, STAT3, Akt, and STAT5 contributing above average (VIP scores greater than 1) to the ADτ, whereas Erk/MAPK ½ and MEK1 to ADτ (**Fig. 4b**).

**Figure 4.**
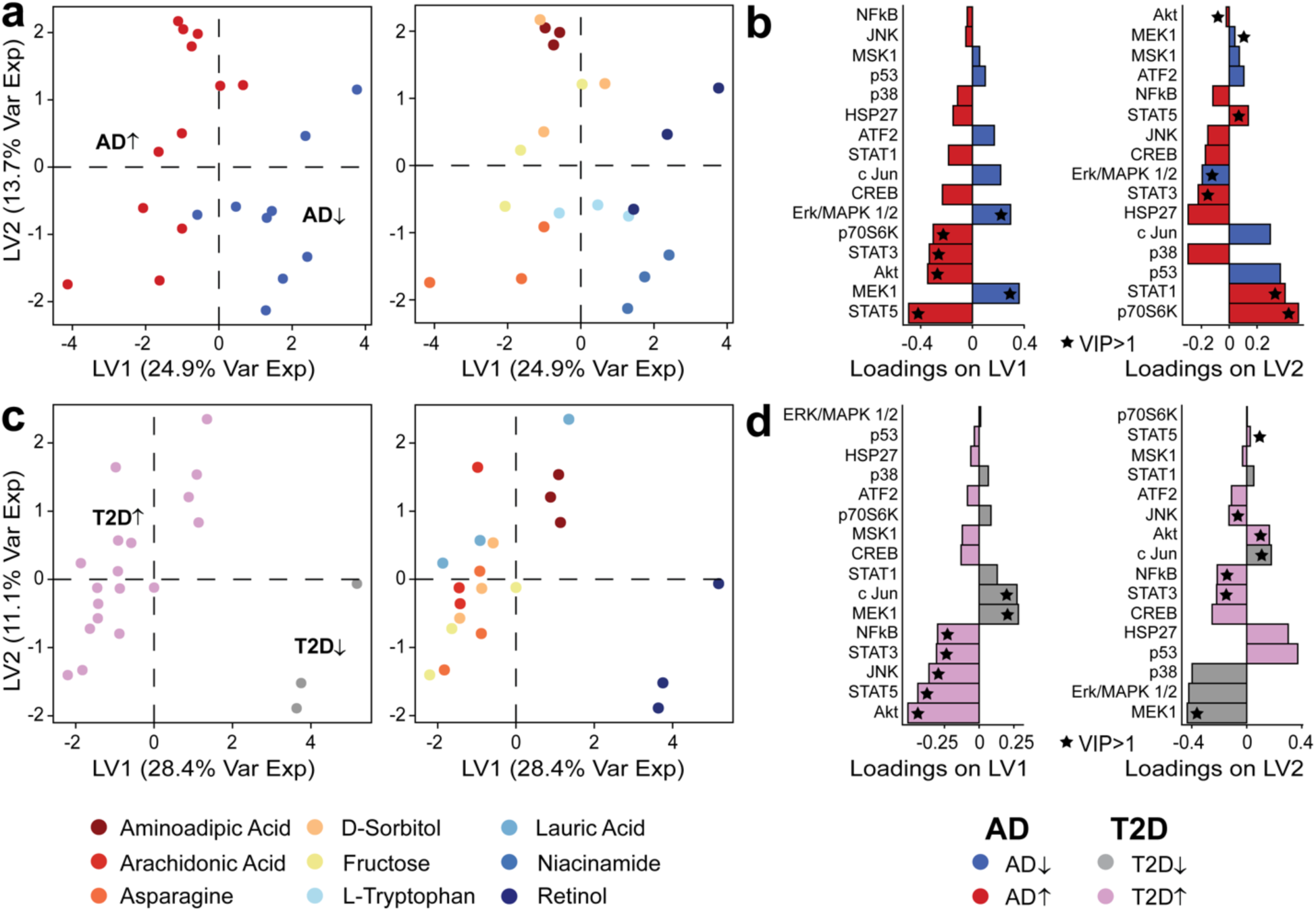
Intracellular protein levels are differentially abundant across metabolites classified by AD or T2D associations. **(a)** Model of AD-protective against AD-associated metabolites**. (b)** The loadings for latent variables 1 and 2 for the AD PLS-DA model **(c)** Model of T2D-protective against T2D-associated classifications. **(d)** The loadings for latent variables 1 and 2 for the T2D PLS-DA model. Loadings for latent variables 1 and 2 denoted with a star indicate a VIP score greater than 1. The color marked on each variable in the loadings represents the analyte with the highest contribution to a metabolite condition.

We also discriminated intracellular signaling levels between metabolites associated with T2D and T2D-protective conditions on LV1 (**Fig. 4c**). In particular, retinol induced a different intracellular neuronal response compared to aminoadipic acid, arachidonic acid, asparagine, D-sorbitol, fructose-6-phosphate, and lauric acid. On T2D LV1, we found NFkB, STAT3, JNK, STAT5, and Akt with a VIP > 1 with its highest contribution to T2Dτ (**Fig. 4d**). These five intracellular proteins are particularly involved in T2D conditions. Oppositely, MEK1 and c-Jun were among the proteins with a VIP score greater than that was associated with T2Dτ.

Among these intracellular signaling proteins, we sought to find pathways that were consistent with a disease direction by synthesizing our findings from all three PLS-DA models. We found MEK1, which is important for regulating cell proliferation and differentiation, to be contributing to AD/T2Dτ on LV1 and LV2 of the four-way model, and to ADτ and the T2Dτ on the AD and T2D PLS-DA models, respectively. This may indicate that MEK1 has an association with protective properties. On the other hand, STAT3, STAT5, and Akt were all proteins that contributed to the AD/T2Dτ classification on the four-way model (both LV1 and LV2) as well as ADτ and T2Dτ on the separate AD and T2D models. The STAT pathways are involved in immune responses, whereas Akt promotes cellular functions such as proliferation and apoptosis. Lastly, we found intracellular proteins with single-disease associations, such as JNK and c-Jun, which were contributing to T2Dτ and T2Dτ, respectively. These two proteins were also supported by the four-way and T2D PLS-DA model.

### Akt and STAT5 are associated with IL-9 signaling, whereas c-Jun and MEK1 are associated with MCP-1 signaling

Having revealed intracellular pathways contributing to the separation among the three PLS-DA models, our next goal was to understand how intracellular and intercellular signaling networks were regulated by metabolites associated with T2D and AD status. To understand this relationship, we integrated our cytokine dataset, which was derived from the same cell culture experiment and wells, with our intracellular protein data to understand the relationship between such signaling molecules.

To do so, we used canonical correlation analysis (CCA) to model links between all intracellular signaling proteins and the two cytokines we previously reported to have differential responses to the metabolite conditions: MCP-1 (monocyte chemoattractant protein-1) and IL-9 (interleukin-9)^8^ (**Fig. 5a**). We previously reported MCP-1, a signaling molecule responsible for monocyte recruitment, to be associated with protective properties of AD and T2D when upregulated and found IL-9, which was associated with promoting mast cell growth, to be related to T2D conditions.

**Figure 5.**
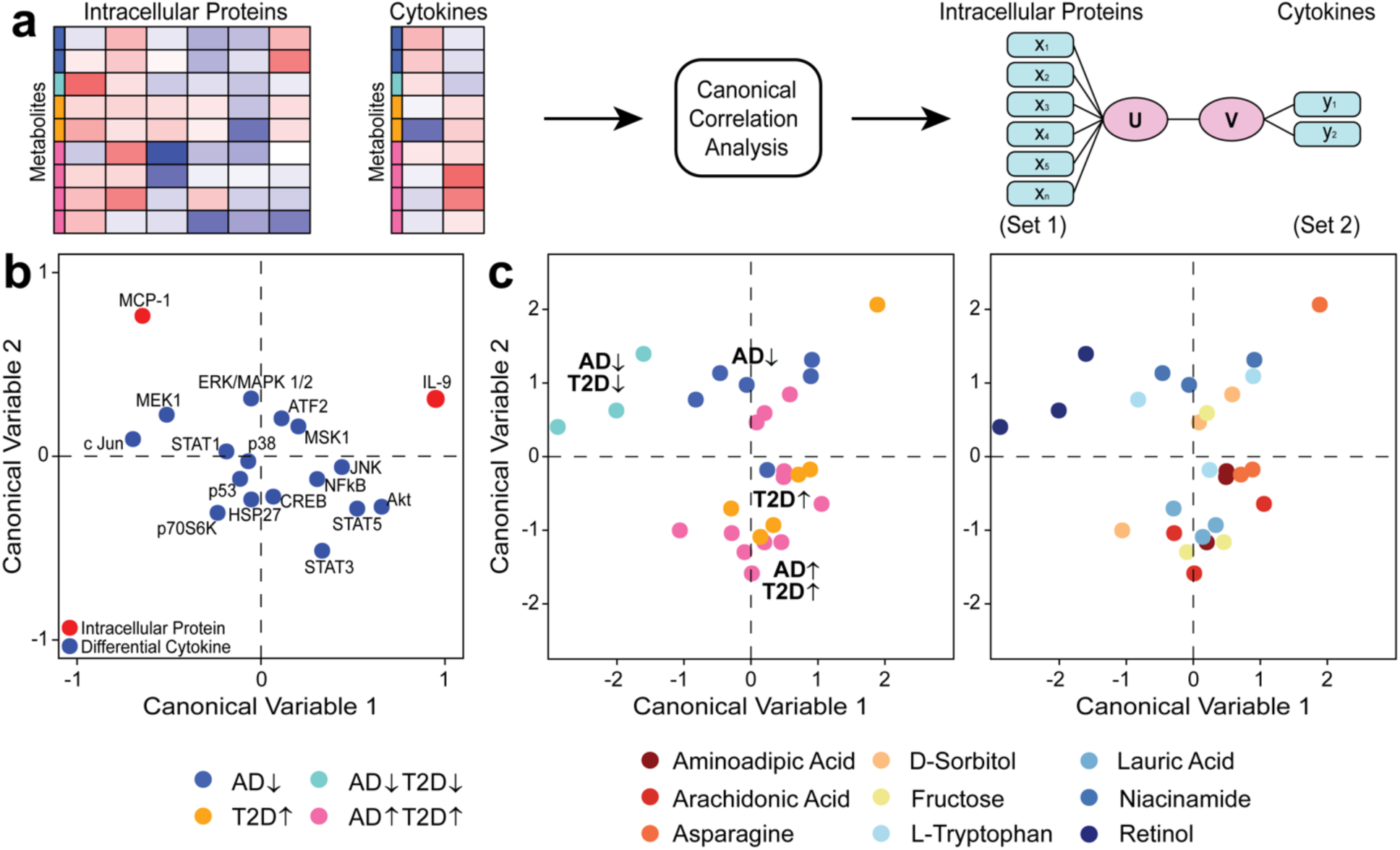
Canonical correlation analysis of intracellular proteins and cytokines. **(a)** All intracellular proteins were used in set 1 for canonical correlation with two cytokines (MCP-1 and IL-9) in set 2. The two cytokines were previously identified to have differential responses to AD- and T2D-associated metabolites. **(b)** Canonical correlation biplot showing the canonical loadings on CV1 and CV2 for both intracellular and cytokine analytes. **(c)** Metabolite conditions were represented by their canonical scores on the first two CVs. Points were annotated by disease condition and respective metabolites.

We visualized the first two canonical variables (CVs) to describe the canonical loadings for the intracellular data and cytokine data. We found that MCP-1 and IL-9 had opposite directionality scores on the first CV (**Fig. 5b**). Along the CV1, we identified intracellular proteins with a negative canonical score (c-Jun, MEK1, STAT1, ERK/MAPK ½, p70S6K, p53, p38, and HSP27) correlated with MCP-1 abundance, while a positive canonical score (CREB, ATF2, MSK1, NFkB, STAT3, JNK, STAT5, and Akt) correlated with IL-9. After visualizing the correlations among intracellular signaling proteins and cytokines, we constructed a scatter plot of the metabolite condition groups to their canonical scores (**Fig. 5c**). We found metabolites with disease-protective properties leaned towards the negative scores on CV1, whereas metabolites associated with disease on the positive end of the spectrum. This finding reinforced our previous findings with retinol’s protective-associated influence with MCP-1.

Using CCA information generated from the IBM SPSS software, we constructed a network connecting the correlation between intracellular proteins and the two cytokines (**Fig. 6**). From our analysis we, found the factors U and V, which each explain the linear combinations for the intracellular and cytokine data respectively, had a canonical correlation of 0.979. (*p* value < 0.001). This correlation represented a strong relationship between the linear combination of the intracellular and cytokine analytes with an 83.9% proportion of variance explained.

**Figure 6.**
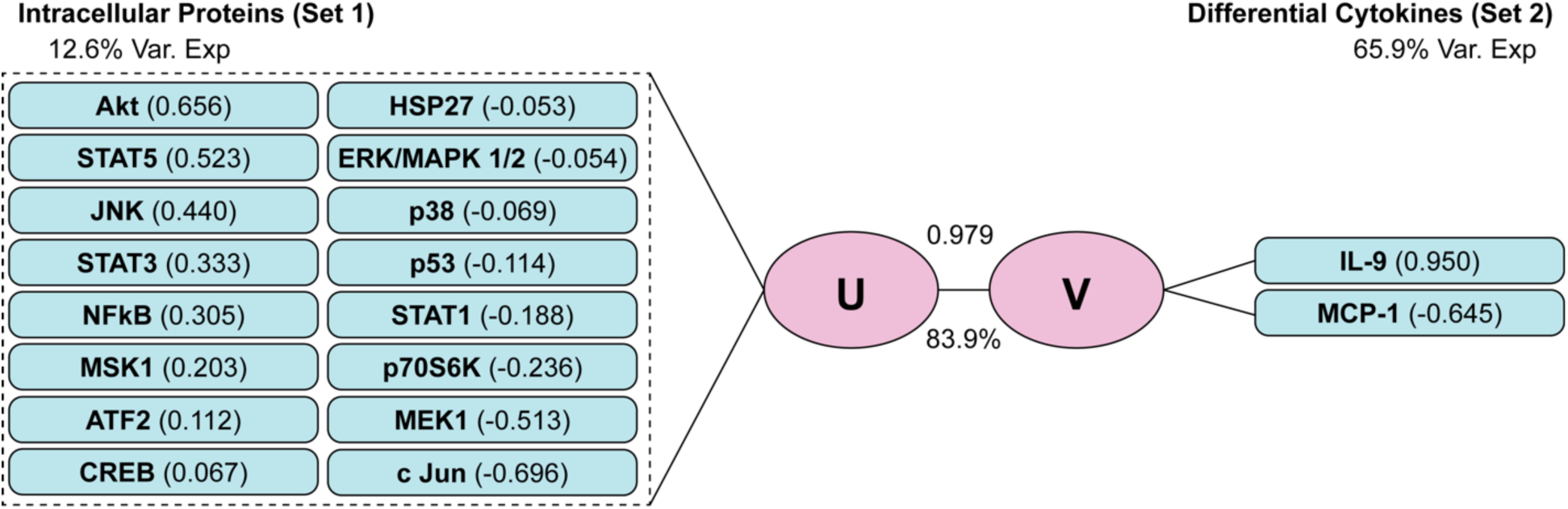
Summarized correlations across intracellular protein and cytokine responses to metabolites associated with T2D and AD conditions. Correlation comparison between intracellular proteins and differential cytokines (IL-9 and MCP-1) on the canonical loadings of the first CV.

We first found that IL-9 and MCP-1 had opposite correlations based on their respective loadings on CV1, supporting our previously tested study that the two cytokines had T2D-associated and AD/T2D-protective properties, respectively. We also identified associations between the intracellular signaling proteins and the cytokines by their direction of magnitude compared across set 1 and set 2 analytes. Of the eight intracellular proteins with a positive correlation, we found Akt and STAT5 were the strongest correlations, with values of 0.656 and 0.523, respectively. We also identified intracellular proteins that had the strongest correlation with MCP-1, which included c-Jun (−0.696) and MEK1 (−0.513).

## DISCUSSION

Intracellular signaling molecules serve a vital role in cell-to-cell communication. In this study, we leveraged experimental and computational approaches to further understand the neuroinflammatory network in response to metabolites associated with T2D and AD. We used an *in vitro* neuronal culture model to quantify different intracellular proteins and cytokines released by neurons in response to different metabolites capable of crossing the BBB into the brain. Our computational analysis identified a cluster of intracellular signaling groups that may have associations with our previously identified MCP-1 and IL-9 cytokines that were differentially abundant in the presence of the same metabolites.

We identified intracellular signaling proteins that were associated with metabolites of disease associations using PLS-DA. Among the intracellular signaling analytes, we found STAT5 and Akt associated with AD and T2D. In a memory function study with mice, Furigo and colleagues found STAT5 knock-out groups had reduced insulin-like growth factor-1 expression in the hippocampus and hypothalamus, showing STAT5 involvement in downstream memory function^48^. STAT5 was also found to be activated under diabetic conditions through low-density lipoprotein glycation and the src kinase pathway^49^. Likewise, the Akt signaling mechanism in both T2D^50^ and AD^51^ is disrupted by insulin resistance, resulting in subsequent alterations to the Akt pathway. For Akt, others have considered targeting the glycogen synthase kinase 3, which is responsible for the phosphorylation of tau and deposition of amyloid beta^52^. The JNK pathway is associated with T2D through insulin resistance and decreased compensatory insulin secretion in different organs including the brain^53^. Interestingly, our modeling did not identify JNK to contribute to AD, despite current efforts investigating this pathway for potential therapeutics^54^. One potential possibility is that JNK was reported to be driven by amyloid beta and tau pathology^55^, which we did not include in our stimulation assay, thus the possible lack of JNK activated by such proteins.

Among our quantified intracellular proteins, we found c-Jun and MEK1 associated with AD- and T2D-protective properties. Increased levels of c-Jun were present in post-mortem brain tissue and found with neurofibrillary tau tangles^56^. Similarly, a group that studied diabetic rats with exposure to high doses of glucose demonstrated high phosphorylated c-Jun expression and neuronal cell death^57^. Thus, lowering c-Jun levels may be of interest for potential therapeutic avenues. Among MEK1 pathways, inhibition of the pathway rescued neurodegeneration in 5XFAD mouse models^58^. In the T2D context, the inhibition of MEK1 improved insulin resistance in the liver of db/db mouse models as well^59^. These relationships we identified from our PLS-DA model also corroborated with our CCA outcomes based on pathways and the disease associations we found on MCP-1 and IL-9.

Using CCA, we mapped a relationship between intracellular signaling molecules with MCP-1 and IL-9 cytokines. In our work, we directly correlated STAT5 and Akt to IL-9 secretion. Likewise, we correlated c-Jun and MEK1 to MCP-1 signaling. Demoulin *et al*. reported STAT5 to be an important mediator for IL-9 function, which was found in the context of lymphoid cells^60^. The Akt pathway was also found to be associated with IL-9 signaling. In the context of synovial T cells in arthritis, the IL-9 cytokine was found to be dependent on PI3K, Akt, and mTOR signaling pathways to induce the proliferation of T cells^61^. Although not directly in the brain, these immune signaling relationships may also be present in the brain.

Similar to IL-9, we found intracellular signaling pathways that are associated with MCP-1, which we previously found to be upregulated in the presence of metabolites associated with AD- or T2D-protective properties. MEK1 signaling was one of many pathways shown to be activated by MCP-1 to regulate vascular endothelial growth factor (VEGF)^62,63^. Interestingly, there is varying evidence on the role of VEGFs in AD, such that in some cases, these growth factors can elicit neuroprotective properties through VEGF ligand and receptor genes^64^. Others showed VEGFs to be elevated in the plasma^65^ of individuals with AD, with additional reports finding increased BBB permeability^66^. However, these differences may be due to confounding variables such as the timeline of the disease and specific tissue samples that were studied. Lastly, we found connections between c-Jun and MCP-1. In a study investigating the use of chemotherapeutic drug treatment for multiple myeloma and neoplasms, they found that inhibition of c-Jun signaling prevented upregulation of MCP-1 in rat models treated with Bortezomib^67^. However, further studies should be conducted to fully understand the cell-to-cell signaling mechanisms in the context of T2D and AD.

Despite our findings, there are limitations and assumptions to our model. Our study focused on a singular communication direction released from primary neuron cells and did not include other important cells such as microglia or astrocytes. Although we excluded other major cell types in the brain, evidence suggests that although these glial cells are actively present in neuroinflammatory pathways, neurons are also capable of releasing neuroinflammatory molecules^68–71^ in both AD^72^ and T2D^73^ conditions. Additionally, our study focused on a metabolite concentration of 300 μM which was established by a thorough literature search and confirmation by a cell viability assay in our previous analysis^8^, which may physiologically differ in the contexts of human translation or other cells. Therefore, prior studies that report differing conclusions may be due to differences in localized tissue samples, animal models, or timeline of disease conditions.

Additionally, our model assumed perfect distinguishment regarding metabolites and their associations to AD and T2D. Despite this label, biological molecules and functions are complex and are difficult to fully associate with a specific condition. Likewise, signaling molecules cannot be fully linked to a disease association since each pathway may serve different cellular communication purposes based on external stimuli. Our findings highlight the correlation between intracellular and cytokine signaling derived from the same biological samples and technical replicates but may not fully map the entire pathway by a direct physical mechanism. Therefore, future experimental considerations may include a variety of other cell types, metabolites, and the use of other cell signaling panels to understand the neuroinflammatory pathway.

Taken together, our investigation revealed differential intracellular signaling responses made by neuronal cells in the presence of metabolites with different disease affiliations. Our computational analysis revealed signaling molecules that are associated with T2D and AD. We also found direct correlational relationships between intracellular signaling proteins and two cytokines (MCP-1 and IL-9) that we previously found differentiated to T2D and AD. Further comprehension of these signaling pathways will improve our understanding of the complex relationship between T2D and AD.

## MATERIALS AND METHODS

### Metabolite selection criteria

We analyzed the same metabolites that were selected in our previous study focused on neuronal cytokine responses^8^. We established an inclusion criterion for candidate metabolites such that the metabolites were differentially abundant in AD or T2D conditions, the metabolite had the potential to cross the BBB, and two or more published studies also reported similar disease associations. We ensured that all metabolites were BBB-permeable^74–82^, and studies with contradictory results for either disease association or BBB-permeability were excluded from our study.

### Animal use and ethics approval

We performed all animal handling and procedures in strict accordance with guidelines approved by the Penn State College of Medicine Institutional Animal Care and Use Committee under protocol number PROTO201800449. This study is also reported in accordance with ARRIVE guidelines.

### Primary neuronal culture

In this study, we used primary neuronal cells derived from embryonic litters. We sacrificed two pregnant CD-1 mice (Charles River Laboratories, strain 022) at gestational day 17 by decapitation using a guillotine. We used a total of 25 embryonic pups shared between the two pregnant mice and sacrificed them using surgical scissors. We placed all embryonic brains in cold HEPES-buffered Hank’s Balanced Salt solution, where we removed the meninges and isolated the cortical cap. We followed an established primary neuronal culture protocol^72^ for this study. Using this method, our team isolated cortices into a conical tube containing a warm embryonic plating medium consisting of Neurobasal Plus (Gibco), 10% fetal bovine serum (Gibco), 1x GlutaMAX (Gibco), and 1x penicillin-streptomycin (10,000 U/mL, Gibco). With a pipette, we manually triturated the embryonic mouse cortices in the embryonic medium and determined the total cell concentration by the Countess II cell counter (Invitrogen). We plated the 6-well plates (coated with 0.1 mg/mL poly-D-lysine) with cell suspension at 5,196,500 cells/well.

After 24 h post-plating, we replaced the embryonic media with neuronal media, which was comprised of Neurobasal Plus (Gibco), 1x B27 Plus supplement (Gibco), 1x GlutaMAX (Gibco), 1x penicillin-streptomycin (10,000 U/mL, Gibco). After 4 days, we replaced half of the neuronal media. On the eighth day post-plating, we inspected the neuronal cells for confluency and the presence of contaminants. Next, we stimulated the neuronal cells with the selected metabolites dissolved in their respective vehicles and media with a final concentration of 300 μM.

After 72 h from stimulating the neuronal cells with metabolites, we collected all cell media and cell lysates for analysis. We obtained cell media and snap-froze each sample in liquid nitrogen for future cytokine quantification. We then collected cell lysates by first carefully washing the 6-well plates with 1X phosphate-buffered saline, adding lysis buffer (Millipore) to the plates, and placing the covered plates on the shaker at 1000 cycles/min for 1 min. The cell solutions were placed into an Eppendorf tube and vortexed for 1 min and placed on ice. After all lysis samples were prepared, they were rested on ice for 20 min, then centrifuged at 5000 x g for 5 min at room temperature. After centrifugation, we carefully removed the supernatants, equally separated the remaining pellets into two Eppendorf tubes, and immediately snap-frozen the samples in liquid nitrogen for future intracellular protein analysis.

### Established metabolite stimulation concentration

We used a metabolite treatment concentration of 300 μM after a previously conducted dose-response viability (Invitrogen, L3224) for various metabolite treatment concentrations^8^. Our previously tested concentrations (1 nM, 3 nM, 10 nM, 10 nM, 30 nM, 100 nM, 300 nM, 1 uM, 3 μM, 10 μM, 30 μM, 100 μM, and 300 μM) were based on independent studies using relevant metabolites on different cell lines^83–91^. Following the primary neuronal culturing protocol, we sacrificed a pregnant CD-1 mouse (gestational day 15) that contained 12 embryos for cell plating in a 96-well plate (density of 175,000 cells/well). We also loaded vehicle control groups that matched each respective metabolite’s solvent. For metabolites requiring dimethyl sulfoxide for dissolving, we ensured the final concentration did not exceed 0.3% for cell viability.

We evaluated neuronal viability using live/dead staining (calcein acetoxymethyl and ethidium homodimer) (Invitrogen, catalog no. L3224). We measured the fluorescence using the SpectraMax i3x plate reader (Molecular Devices) and reported no significant decline in cell viability with increasing concentrations. As a result, we established 300 μM for our concentration based on an assumption that high acute exposure to a treatment condition will mimic a neuronal response to low, chronic exposure.

### Bicinchoninic acid assay for protein quantification

We performed a bicinchoninic acid (BCA) assay to standardize protein concentrations loaded into each well of the 384-plate for the Luminex assay. We quantified the total protein content from the lysed neuronal cell samples using the Pierce™ BCA assay kit (Fisher, 23225) according to the manufacturer’s manual. We loaded 10 μL of each sample, along with 10 μL of standards into 96-well plates in triplicates, and a total of 200 μL of working solution from the BCA assay kit to each well. We covered the plates with an adhesive seal and placed them on a shaker at 400 cycles/min for 30 s, then in an incubator for 30 min. Using the SpectraMax i3x plate reader (Molecular Devices), we recorded the absorbances for each standard and sample. We then quantified sample concentrations by linear regression using the standards from each respective plate.

### Intracellular signaling protein quantification

We quantified intracellular signaling proteins using the Luminex FLEXMAP 3D using the Milliplex 10-plex MAPK/SAPK signaling magnetic bead kit (Millipore, 48-660MAG) and Milliplex 9-plex multi-pathway magnetic bead kit (Millipore, 48-680MAG). The MAPK/SAPK signaling magnetic bead kit detects ATF2, JNK, HSP27, p38, ERK/MAPK ½, p53, MEK1, MSK1, STAT1, and c-Jun. The multi-pathway magnetic bead kit detects ERK/MAPK ½, Akt, STAT3, p70S6K,

NFkB, STAT5, CREB, and p38. We modified the manufacturer’s protocol to accommodate a 384-well format by adding magnetic beads and antibodies at a reduced volume to each kit as demonstrated by other studies^8,92,93^. We loaded 10 μg of protein lysate, along with blanks and quality controls into the 384-well plate in technical triplicate for the Luminex assay. We performed instrument calibration and verification prior to running the Luminex assay.

### Cytokine signaling quantification

From a previous study^8^, we quantified the intensities of cytokines on the Luminex Bio-Plex 3D platform. We used the Bio-Rad BioPlex 23-plex pro mouse cytokine kit (Bio-Rad, M60009RDPD). The kit contains a panel to quantify levels of cytokines, included eotaxin, G-CSF, IFN-γ, IL-17A, IL-1β, IL-1α, IL-10, IL-12p40, IL-12p70, IL-13, IL-2, IL-3, IL-4, IL-5, IL-6, IL-9, KC, MCP-1, MIP-1β, MIP-1α, RANTES, and TNF-α. We modified the manufacturer’s protocol to accommodate a 384-well format by adding magnetic beads and antibodies at a reduced volume^92–94^. We loaded samples, blanks, and quality control standards into a 384-well plate in technical triplicate and quantified each cytokine. We also performed instrument calibration and verification before running the Luminex assay.

### Intracellular protein data pre-processing

We performed all data pre-processing in R. Before combining the intracellular kits, we performed a principal component analysis to ensure there were no visual batch effects by separation (**Supplementary Fig. S1**), although the assays were run on the same machine, from the same biological animals, and the same technical replicates. The principal component analysis offers a visualization approach to compare potential differences across the batches.

After verifying that there were no batch effects, we averaged any duplicate intracellular signaling pathways shared across the two kits to prevent bias from excess readings from the same pathway. We then transformed the data by taking the log_2_ ratio of the individual measurement by the average of their respective triplicate vehicle intracellular protein concentrations. No intracellular protein data was excluded from the analysis.

### Cytokine data pre-processing

From our previous study, we excluded cytokines that contained more than 25% of readings below the lower limit of quantification. As a result, we excluded eight cytokines (G-CSF, IL-2, IL-3, IL-4, IL-5, IL13, IL-17A, and TNF-α) from downstream analysis. Any data for the other fifteen remaining cytokines were processed by substituting values that were below the lowest limit of quantification with the lowest standard value for that cytokine (2.1% of the data). Afterward, we transformed the data by taking the log_2_ ratio of the individual measurements by the average of their respective triplicate vehicle concentrations.

### Hierarchical clustering

We hierarchically clustered intracellular signaling proteins to identify any relative changes with respect to the different metabolite groups. We applied a z-score transformation on the log_2_FC intracellular protein dataset before constructing the heatmap. To visualize the heatmap, we used the *pheatmap*^95^ package (ver. 1.0.12) in R.

### Partial least squares discriminant analysis

To further understand the patterns and signaling relationships of multiple intracellular proteins simultaneously, we performed PLS-DA, a multivariate dimensional-reduction approach in R. Using the *mixOmics*^96^ package (ver. 6.26.0), we constructed three separate PLS-DA models with different metabolite classifications to better understand the relationships the neuronal intracellular signaling responses may have to stimulation. The three models included 4-way discrimination across all metabolite groups (ADτ, T2Dτ, AD/T2Dτ, and AD/T2Dτ), a model focused on AD-associated metabolites (ADτ vs ADτ), and a model focused on T2D-associated metabolites (T2Dτ vs T2Dτ). We then interpreted the first two latent variables from each respective PLS-DA model.

We identified the most important intracellular proteins contributing to the separation across the classification groups by using the calculated VIP score, which represents the strength of contribution from each protein to the PLS-DA results. To determine respective VIP scores for each intracellular protein, we used the *mixOmics*^96^ package (ver. 6.26.0). We defined a VIP score threshold greater than 1 to contribute to the PLS-DA model above average.

### Canonical correlation analysis

To correlate the intracellular signaling proteins to the cytokine responses made by the neuronal cells, we used the IBM SPSS Statistics (ver. 30.0.0.0) software to perform a CCA. We combined the log_2_FC of the respective cytokine and intracellular signaling datasets by columns and matched their respective rows (metabolite stimulation groups and wells) into a singular dataset. We applied the “Canonical Correlation” analysis function and defined the intracellular pathways (Akt, ATF2, c-Jun, CREB, Erk/MAKP ½, HSP27, JNK, MEK1, MSK1, NFkB, p38, p53, p70S6K, STAT1, STAT3, and STAT5) in set 1. For set 2, we defined the cytokine signals MCP-1 and IL-9. For data visualization, we used the *CCA*^97^ (ver. 1.2.2) package in R to generate scatterplots of the canonical loadings.

### Statistical Analysis

We used a Kruskal-Wallis test adjusted by Benjamini-Hochberg to determine the statistical significance of intracellular protein levels across metabolites separated by their treatment association (ADτ, T2Dτ, AD/T2Dτ, and AD/T2Dτ). We defined an FDR *q* value less than 0.10 to be significant. For any proteins that were found significant from the Kruskal-Wallis test, we performed a Mann-Whitney test to identify the specific responses that were significant across the four established conditions. In this case, we set a *p* value less than 0.05 to be significant.

## ABBREVIATIONS

AD: Alzheimer’s disease
Akt: Protein kinase B
BBB: Blood-brain barrier
BCA: Bicinchoninic acid
CCA: Canonical correlation analysis
CV: Canonical variable
FDR: False discovery rate
IL-9: Interleukin-9
JNK: c-Jun N-terminal kinase
LV: Latent variable
MEK1: mitogen-activated protein kinase 1
MCP-1: Monocyte chemoattractant protein-1
PLS-DA: Partial least squares discriminant analysis
STAT1: Signal transducer and activator transcription 1
STAT5: Signal transducer and activator transcription 5
T2D: Type 2 diabetes
VEGF: Vascular endothelial growth factor
VIP: Variable importance in projection

## ACKNOWLEDGEMENTS

The research work in this study is supported by an award from the Good Ventures Foundation, Open Philanthropy, and start-up funds from Purdue University Weldon School of Biomedical Engineering (DKB and BKB). This work is also supported by R21AG068532 from the National Institute on Aging (EAP). BKB is supported by the NIH T32 predoctoral fellowship (T32DK101001) from the National Institute of Diabetes and Digestive and Kidney Diseases. BKB also acknowledges the National Science Foundation for support under the Graduate Research Fellowship Program (DGE-1842166). MKK is supported by the NIH T32 training fellowship from the National Institute of Neurological Disorders and Stroke (T32NS115667). RMFB is supported by the NIH NRSA predoctoral fellowship from the National Institute on Aging (F31AG071131). The authors also appreciate Javier Munoz-Briones for his support in the cytokine Luminex assay.

## AUTHOR CONTRIBUTIONS

**BKB:** Conceptualization, data curation, formal analysis, funding acquisition, investigation, methodology, visualization, writing-original draft, writing-review & editing. **MKK:** Formal analysis, investigation, methodology, writing-review & editing. **RMFB:** Formal analysis, investigation, methodology, writing-review & editing. **EAP:** Conceptualization, funding acquisition, methodology, project administration, resources, writing-review & editing. **DKB:** Conceptualization, funding acquisition, methodology, project administration, resources, writing-review & editing.

## DATA AVAILABILITY

Generated data for analysis is included and available in the **Supplementary Files** of this article.

## CODE AVAILABILITY

All code is publicly available at https://github.com/Brubaker-Lab/AD-T2D-IntracellularSignaling.

## SUPPLEMENTARY INFORMATION

**Supplementary Figure S1.**
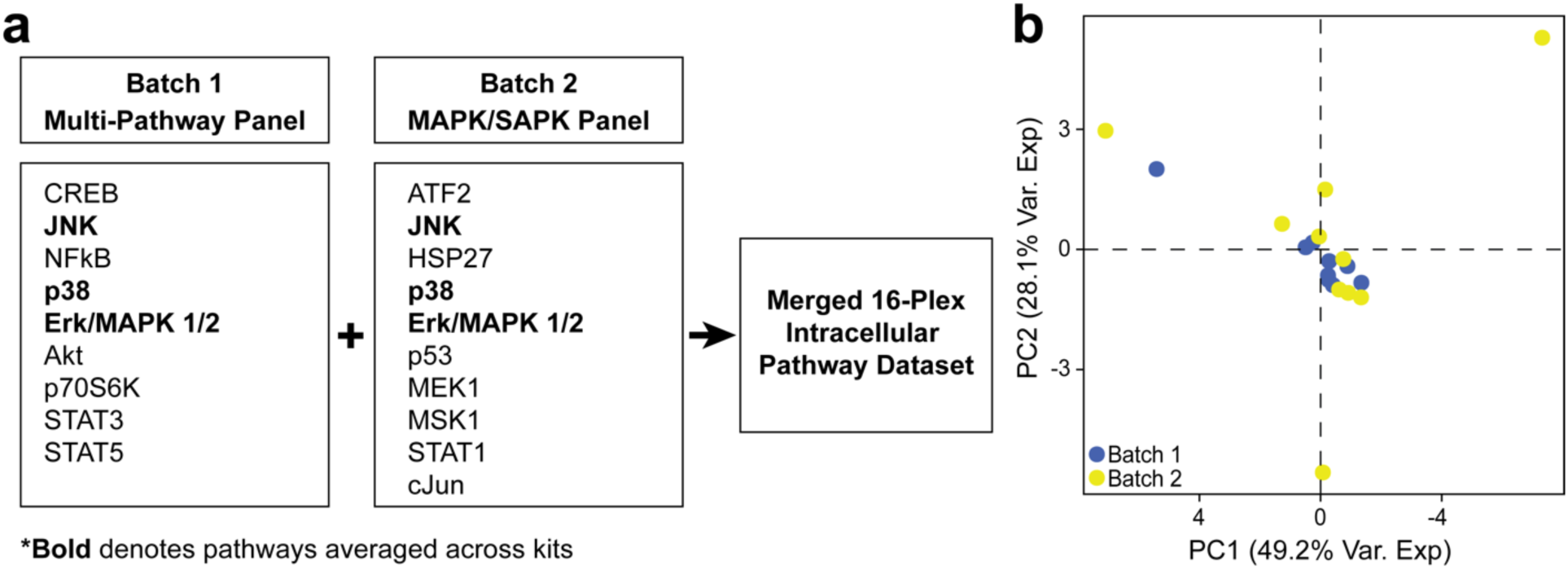
Data merging of quantified intracellular proteins. **(a)** Separate Luminex immunoassays were merged into 16 total analytes, where those repeated twice were averaged. **(b)** Principal component analysis comparing the batches before the averaging of data to confirm no batch effects were present.

**Supplementary Figure S2.**
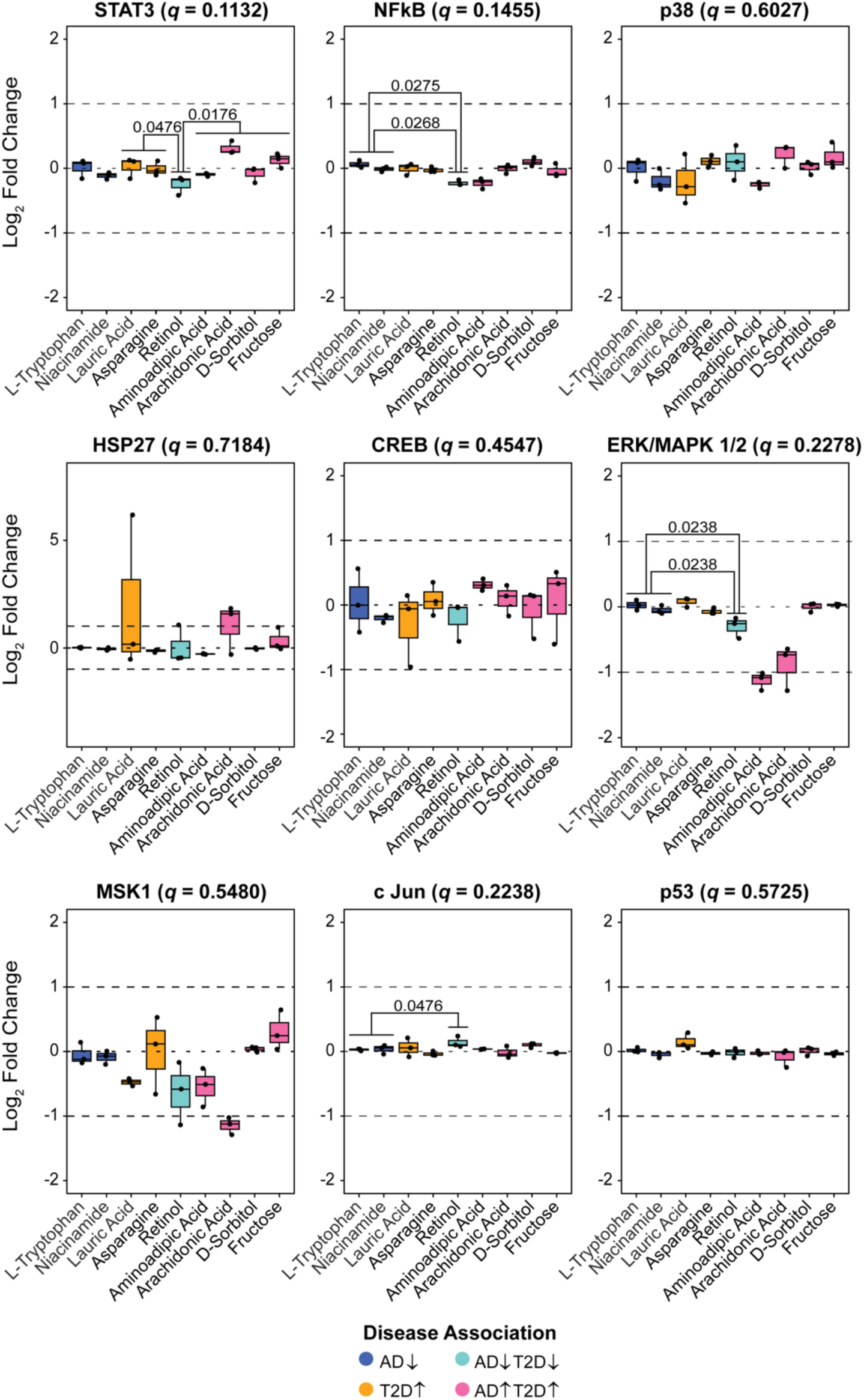
Pairwise testing of non-significant intracellular pathways. All pathways were found to have no significance based on the Kruskal-Wallis test adjusted by the Benjamini-Hochberg method (FDR *q* < 0.10). Comparisons across metabolite conditions were done by a Mann-Whitney test (*p* < 0.05).

## REFERENCES

1. Zilkens, R. R., Davis, W. A., Spilsbury, K., Semmens, J. B. & Bruce, D. G. Earlier Age of Dementia Onset and Shorter Survival Times in Dementia Patients With Diabetes. American Journal of Epidemiology 177, 1246–1254 (2013).

2. Wang, K.-C. et al. Risk of Alzheimer’s Disease in Relation to Diabetes: A Population-Based Cohort Study. Neuroepidemiology 38, 237–244 (2012).

3. Janson, J. et al. Increased Risk of Type 2 Diabetes in Alzheimer Disease. Diabetes 53, 474– 481 (2004).

4. Gudala, K., Bansal, D., Schifano, F. & Bhansali, A. Diabetes mellitus and risk of dementia: A meta-analysis of prospective observational studies. Journal of Diabetes Investigation 4, 640–650 (2013).

5. Cheng, G., Huang, C., Deng, H. & Wang, H. Diabetes as a risk factor for dementia and mild cognitive impairment: a meta-analysis of longitudinal studies. Internal Medicine Journal 42, 484–491 (2012).

6. Srodulski, S. et al. Neuroinflammation and neurologic deficits in diabetes linked to brain accumulation of amylin. Molecular Neurodegeneration 9, 30 (2014).

7. Paul, S., Bhardwaj, J. & Binukumar, B. K. Cdk5-mediated oligodendrocyte myelin breakdown and neuroinflammation: Implications for the link between Type 2 Diabetes and Alzheimer’s disease. Biochimica et Biophysica Acta (BBA) -Molecular Basis of Disease 1870, 166986 (2024).

8. Ball, B. K., Kuhn, M. K., Fleeman Bechtel, R. M., Proctor, E. A. & Brubaker, D. K. Differential responses of primary neuron-secreted MCP-1 and IL-9 to type 2 diabetes and Alzheimer’s disease-associated metabolites. Sci Rep 14, 12743 (2024).

9. Ball, B. K., Proctor, E. A. & Brubaker, D. K. Cross-Species Modeling Identifies Gene Signatures in Type 2 Diabetes Mouse Models Predictive of Inflammatory and Estrogen Signaling Pathways Associated with Alzheimer’s Disease Outcomes in Humans. In Biocomputing 2025 426–440 (WORLD SCIENTIFIC, 2024). doi:10.1142/9789819807024_0031.

10. Ball, B. K., Park, J. H., Proctor, E. A. & Brubaker, D. K. Cross-disease modeling of peripheral blood identifies biomarkers of type 2 diabetes predictive of Alzheimer’s disease. 2024.12.11.627991 Preprint at 10.1101/2024.12.11.627991 (2024).

11. Huang, C., Luo, J., Wen, X. & Li, K. Linking Diabetes Mellitus with Alzheimer’s Disease: Bioinformatics Analysis for the Potential Pathways and Characteristic Genes. Biochem Genet 60, 1049–1075 (2022).

12. Chung, Y. & Lee, H. Correlation between Alzheimer’s disease and type 2 diabetes using non-negative matrix factorization. Sci Rep 11, 15265 (2021).

13. Chowdhury, U. N., Islam, M. B., Ahmad, S. & Moni, M. A. Network-based identification of genetic factors in ageing, lifestyle and type 2 diabetes that influence to the progression of Alzheimer’s disease. Informatics in Medicine Unlocked 19, 100309 (2020).

14. Kadry, H., Noorani, B. & Cucullo, L. A blood–brain barrier overview on structure, function, impairment, and biomarkers of integrity. Fluids and Barriers of the CNS 17, 69 (2020).

15. Porkoláb, G. et al. Synergistic induction of blood–brain barrier properties. Proceedings of the National Academy of Sciences 121, e2316006121 (2024).

16. Hawkins, B. T., Lundeen, T. F., Norwood, K. M., Brooks, H. L. & Egleton, R. D. Increased blood–brain barrier permeability and altered tight junctions in experimental diabetes in the rat: contribution of hyperglycaemia and matrix metalloproteinases. Diabetologia 50, 202–211 (2007).

17. Sengillo, J. D. et al. Deficiency in Mural Vascular Cells Coincides with Blood–Brain Barrier Disruption in Alzheimer’s Disease. Brain Pathology 23, 303–310 (2013).

18. Zenaro, E. et al. Neutrophils promote Alzheimer’s disease–like pathology and cognitive decline via LFA-1 integrin. Nat Med 21, 880–886 (2015).

19. Ono, K. & Yamada, M. Vitamin A and Alzheimer’s disease. Geriatrics & Gerontology International 12, 180–188 (2012).

20. Davidson, T. L. et al. The effects of a high-energy diet on hippocampal-dependent discrimination performance and blood–brain barrier integrity differ for diet-induced obese and diet-resistant rats. Physiology & Behavior 107, 26–33 (2012).

21. Janelidze, S. et al. Increased blood-brain barrier permeability is associated with dementia and diabetes but not amyloid pathology or APOE genotype. Neurobiology of Aging 51, 104– 112 (2017).

22. Sandireddy, R., Yerra, V. G., Areti, A., Komirishetty, P. & Kumar, A. Neuroinflammation and Oxidative Stress in Diabetic Neuropathy: Futuristic Strategies Based on These Targets. International Journal of Endocrinology 2014, 674987 (2014).

23. Heneka, M. T. et al. Neuroinflammation in Alzheimer’s disease. The Lancet Neurology 14, 388–405 (2015).

24. Lampron, A., ElAli, A. & Rivest, S. Innate Immunity in the CNS: Redefining the Relationship between the CNS and Its Environment. Neuron 78, 214–232 (2013).

25. Xanthos, D. N. & Sandkühler, J. Neurogenic neuroinflammation: inflammatory CNS reactions in response to neuronal activity. Nat Rev Neurosci 15, 43–53 (2014).

26. West, A. E., Griffith, E. C. & Greenberg, M. E. Regulation of transcription factors by neuronal activity. Nat Rev Neurosci 3, 921–931 (2002).

27. Pegtel, D. M., Peferoen, L. & Amor, S. Extracellular vesicles as modulators of cell-to-cell communication in the healthy and diseased brain. Philosophical Transactions of the Royal Society B: Biological Sciences 369, 20130516 (2014).

28. Weisman, D., Hakimian, E. & Ho, G. J. Interleukins, Inflammation, and Mechanisms of Alzheimer’s Disease. in Vitamins & Hormones vol. 74 505–530 (Academic Press, 2006).

29. Domingues, C., A.B. da Cruz e Silva, O. & Henriques, A. Impact of Cytokines and Chemokines on Alzheimer’s Disease Neuropathological Hallmarks. Current Alzheimer Research 14, 870–882 (2017).

30. Zipp, F., Bittner, S. & Schafer, D. P. Cytokines as emerging regulators of central nervous system synapses. Immunity 56, 914–925 (2023).

31. Wang, L.-W., Tu, Y.-F., Huang, C.-C. & Ho, C.-J. JNK signaling is the shared pathway linking neuroinflammation, blood–brain barrier disruption, and oligodendroglial apoptosis in the white matter injury of the immature brain. Journal of Neuroinflammation 9, 175 (2012).

32. Li, X. et al. Downregulation of STAT1 improved learning and memory impairments in aging mice. Neuroscience Letters 850, 138155 (2025).

33. Zhou, W. et al. Longitudinal multi-omics of host–microbe dynamics in prediabetes. Nature 569, 663–671 (2019).

34. Kamoshita, K. et al. Lauric acid impairs insulin-induced Akt phosphorylation by upregulating SELENOP expression via HNF4α induction. American Journal of Physiology-Endocrinology and Metabolism 322, E556–E568 (2022).

35. Luo, H.-H., Feng, X.-F., Yang, X.-L., Hou, R.-Q. & Fang, Z.-Z. Interactive effects of asparagine and aspartate homeostasis with sex and age for the risk of type 2 diabetes risk. Biol Sex Differ 11, 58 (2020).

36. Hunsberger, H. C. et al. Divergence in the metabolome between natural aging and Alzheimer’s disease. Sci Rep 10, 12171 (2020).

37. Ross, A. P., Bartness, T. J., Mielke, J. G. & Parent, M. B. A high fructose diet impairs spatial memory in male rats. Neurobiology of Learning and Memory 92, 410–416 (2009).

38. Balakumar, M. et al. High-fructose diet is as detrimental as high-fat diet in the induction of insulin resistance and diabetes mediated by hepatic/pancreatic endoplasmic reticulum (ER) stress. Mol Cell Biochem 423, 93–104 (2016).

39. Amtul, Z., Uhrig, M., Wang, L., Rozmahel, R. F. & Beyreuther, K. Detrimental effects of arachidonic acid and its metabolites in cellular and mouse models of Alzheimer’s disease: structural insight. Neurobiology of Aging 33, 831.e21–831.e31 (2012).

40. Williams, E. S., Baylin, A. & Campos, H. Adipose tissue arachidonic acid and the metabolic syndrome in Costa Rican adults. Clinical Nutrition 26, 474–482 (2007).

41. Wang, T. J. et al. 2-Aminoadipic acid is a biomarker for diabetes risk. J Clin Invest 123, 4309–4317 (2013).

42. Li, C.-H. et al. Long-term consumption of the sugar substitute sorbitol alters gut microbiome and induces glucose intolerance in mice. Life Sciences 305, 120770 (2022).

43. Green, K. N. et al. Nicotinamide Restores Cognition in Alzheimer’s Disease Transgenic Mice via a Mechanism Involving Sirtuin Inhibition and Selective Reduction of Thr231-Phosphotau. J Neurosci 28, 11500–11510 (2008).

44. Dalmasso, M. C. et al. Nicotinamide as potential biomarker for Alzheimer’s disease: A translational study based on metabolomics. Front Mol Biosci 9, 1067296 (2023).

45. Porter, R. J. et al. Cognitive Deficit Induced by Acute Tryptophan Depletion in Patients With Alzheimer’s Disease. AJP 157, 638–640 (2000).

46. Noristani, H. N., Verkhratsky, A. & Rodríguez, J. J. High tryptophan diet reduces CA1 intraneuronal β-amyloid in the triple transgenic mouse model of Alzheimer’s disease. Aging Cell 11, 810–822 (2012).

47. Dorey, C. K., Gierhart, D., Fitch, K. A., Crandell, I. & Craft, N. E. Low Xanthophylls, Retinol, Lycopene, and Tocopherols in Grey and White Matter of Brains with Alzheimer’s Disease. Journal of Alzheimer’s Disease 94, 1–17 (2023).

48. Furigo, I. C. et al. Brain STAT5 signaling modulates learning and memory formation. Brain Struct Funct 223, 2229–2241 (2018).

49. Brizzi, M. F. et al. STAT5 Activation Induced by Diabetic LDL Depends on LDL Glycation and Occurs Via src Kinase Activity. Diabetes 51, 3311–3317 (2002).

50. Huang, X., Liu, G., Guo, J. & Su, Z. The PI3K/AKT pathway in obesity and type 2 diabetes. Int J Biol Sci 14, 1483–1496 (2018).

51. Limantoro, J., de Liyis, B. G. & Sutedja, J. C. Akt signaling pathway: a potential therapy for Alzheimer’s disease through glycogen synthase kinase 3 beta inhibition. *The Egyptian Journal of Neurology*, Psychiatry and Neurosurgery 59, 147 (2023).

52. Pan, J. et al. The role of PI3K signaling pathway in Alzheimer’s disease. Front. Aging Neurosci. 16, (2024).

53. Yung, J. H. M. & Giacca, A. Role of c-Jun N-terminal Kinase (JNK) in Obesity and Type 2 Diabetes. Cells 9, 706 (2020).

54. Yarza, R., Vela, S., Solas, M. & Ramirez, M. J. c-Jun N-terminal Kinase (JNK) Signaling as a Therapeutic Target for Alzheimer’s Disease. Front Pharmacol 6, 321 (2016).

55. Solas, M. et al. JNK Activation in Alzheimer’s Disease Is Driven by Amyloid β and Is Associated with Tau Pathology. ACS Chem Neurosci 14, 1524–1534 (2023).

56. Pearson, A. G., Byrne, U. T. E., MacGibbon, G. A., Faull, R. L. M. & Dragunow, M. Activated c-Jun is present in neurofibrillary tangles in Alzheimer’s disease brains. Neuroscience Letters 398, 246–250 (2006).

57. Oshitari, T., Bikbova, G. & Yamamoto, S. Increased expression of phosphorylated c-Jun and phosphorylated c-Jun N-terminal kinase associated with neuronal cell death in diabetic and high glucose exposed rat retinas. Brain Research Bulletin 101, 18–25 (2014).

58. Chun, Y. S. et al. MEK1/2 inhibition rescues neurodegeneration by TFEB-mediated activation of autophagic lysosomal function in a model of Alzheimer’s Disease. Mol Psychiatry 27, 4770–4780 (2022).

59. Ueyama, A. et al. Inhibition of MEK1 Signaling Pathway in the Liver Ameliorates Insulin Resistance. J Diabetes Res 2016, 8264830 (2016).

60. Demoulin, J.-B. et al. STAT5 Activation Is Required for Interleukin-9-dependent Growth and Transformation of Lymphoid Cells1. Cancer Research 60, 3971–3977 (2000).

61. Kundu-Raychaudhuri, S., Abria, C. & Raychaudhuri, S. P. IL-9, a local growth factor for synovial T cells in inflammatory arthritis. Cytokine 79, 45–51 (2016).

62. Lien, M.-Y. et al. Monocyte Chemoattractant Protein 1 Promotes VEGF-A Expression in OSCC by Activating ILK and MEK1/2 Signaling and Downregulating miR-29c. Front Oncol 10, 592415 (2020).

63. Zhao, H.-Y. et al. MCP-1 facilitates VEGF production by removing miR-374b-5p blocking of VEGF mRNA translation. Biochemical Pharmacology 206, 115334 (2022).

64. Mahoney, E. R. et al. Brain expression of the vascular endothelial growth factor gene family in cognitive aging and alzheimer’s disease. Mol Psychiatry 26, 888–896 (2021).

65. Chiappelli, M. et al. VEGF Gene and Phenotype Relation with Alzheimer’s Disease and Mild Cognitive Impairment. Rejuvenation Research 9, 485–493 (2006).

66. Miners, J. S., Schulz, I. & Love, S. Differing associations between Aβ accumulation, hypoperfusion, blood–brain barrier dysfunction and loss of PDGFRB pericyte marker in the precuneus and parietal white matter in Alzheimer’s disease. J Cereb Blood Flow Metab 38, 103–115 (2018).

67. Liu, C. et al. Upregulation of CCL2 via ATF3/c-Jun interaction mediated the Bortezomib-induced peripheral neuropathy. Brain, Behavior, and Immunity 53, 96–104 (2016).

68. Rubio-Perez, J. M. & Morillas-Ruiz, J. M. A Review: Inflammatory Process in Alzheimer’s Disease, Role of Cytokines. The Scientific World Journal 2012, 1–15 (2012).

69. Yamamoto, T., Yamashita, A., Yamada, K. & Hata, R.-I. Immunohistochemical localization of chemokine CXCL14 in rat hypothalamic neurons. Neuroscience Letters 487, 335–340 (2011).

70. Tchelingerian, J. L., Vignais, L. & Jacque, C. TNF alpha gene expression is induced in neurones after a hippocampal lesion. Neuroreport 5, 585–588 (1994).

71. Su, J. et al. Cell–cell communication: new insights and clinical implications. Sig Transduct Target Ther 9, 1–52 (2024).

72. Wood, L. B. et al. Identification of neurotoxic cytokines by profiling Alzheimer’s disease tissues and neuron culture viability screening. Sci Rep 5, 16622 (2015).

73. Saleh, A. et al. Tumor necrosis factor-α elevates neurite outgrowth through an NF-κB-dependent pathway in cultured adult sensory neurons: Diminished expression in diabetes may contribute to sensory neuropathy. Brain Research 1423, 87–95 (2011).

74. Aristyani, S., Nur, M. I., Widyarti, S. & Sumitro, S. B. In silico study of active compounds ADMET Profiling in Curcuma xanthorrhiza Roxb and Tamarindus indica as Tuberculosis Treatment. J Jamu Indo 3, 101–108 (2018).

75. Rizzari, C. et al. Asparagine levels in the cerebrospinal fluid of children with acute lymphoblastic leukemia treated with pegylated-asparaginase in the induction phase of the AIEOP-BFM ALL 2009 study. Haematologica 104, 1812–1821 (2019).

76. Page, K. A. et al. Effects of Fructose vs Glucose on Regional Cerebral Blood Flow in Brain Regions Involved With Appetite and Reward Pathways. JAMA 309, 63–70 (2013).

77. Murphy, E. J. Blood–brain barrier and brain fatty acid uptake: Role of arachidonic acid and PGE2. Journal of Neurochemistry 135, 845–848 (2015).

78. Kelly, M., Widjaja-Adhi, M. A. K., Palczewski, G. & von Lintig, J. Transport of vitamin A across blood–tissue barriers is facilitated by STRA6. FASEB J 30, 2985–2995 (2016).

79. Richard, D. M. et al. L-Tryptophan: Basic Metabolic Functions, Behavioral Research and Therapeutic Indications. Int J Tryptophan Res 2, 45–60 (2009).

80. Fricker, R. A., Green, E. L., Jenkins, S. I. & Griffin, S. M. The Influence of Nicotinamide on Health and Disease in the Central Nervous System. Int J□Tryptophan□Res 11, 1178646918776658 (2018).

81. Zaragozá, R. Transport of Amino Acids Across the Blood-Brain Barrier. Front Physiol 11, 973 (2020).

82. Xu, J. et al. Elevation of brain glucose and polyol-pathway intermediates with accompanying brain-copper deficiency in patients with Alzheimer’s disease: metabolic basis for dementia. Sci Rep 6, 27524 (2016).

83. Nonaka, Y. et al. Lauric Acid Stimulates Ketone Body Production in the KT-5 Astrocyte Cell Line. J. Oleo Sci. 65, 693–699 (2016).

84. Krall, A. S., Xu, S., Graeber, T. G., Braas, D. & Christofk, H. R. Asparagine promotes cancer cell proliferation through use as an amino acid exchange factor. Nat Commun 7, 11457 (2016).

85. Lodha, D. & Subramaniam, J. R. High Fructose Negatively Impacts Proliferation of NSC-34 Motor Neuron Cell Line. J Neurosci Rural Pract 13, 114–118 (2022).

86. Chen, Q., Galleano, M. & Cederbaum, A. I. Cytotoxicity and Apoptosis Produced by Arachidonic Acid in Hep G2 Cells Overexpressing Human Cytochrome P4502E1. Journal of Biological Chemistry 272, 14532–14541 (1997).

87. Huck, S., Grass, F. & Hatten, M. E. Gliotoxic effects of α-aminoadipic acid on monolayer cultures of dissociated postnatal mouse cerebellum. Neuroscience 12, 783–791 (1984).

88. Lu, X., Li, C., Wang, Y.-K., Jiang, K. & Gai, X.-D. Sorbitol induces apoptosis of human colorectal cancer cells via p38 MAPK signal transduction. Oncol Lett 7, 1992–1996 (2014).

89. Vesprini, N. D. & Spencer, G. E. Retinoic acid induces changes in electrical properties of adult neurons in a dose- and isomer-dependent manner. J Neurophysiol 111, 1318–1330 (2014).

90. Silk, J. D. et al. IDO Induces Expression of a Novel Tryptophan Transporter in Mouse and Human Tumor Cells. The Journal of Immunology 187, 1617–1625 (2011).

91. Griffin, S. M., Pickard, M. R., Hawkins, C. P., Williams, A. C. & Fricker, R. A. Nicotinamide restricts neural precursor proliferation to enhance catecholaminergic neuronal subtype differentiation from mouse embryonic stem cells. PLoS One 15, e0233477 (2020).

92. Fleeman, R. M., Kuhn, M. K., Chan, D. C. & Proctor, E. A. Apolipoprotein E ε4 modulates astrocyte neuronal support functions in the presence of amyloid-β. Journal of Neurochemistry 165, 536–549 (2023).

93. Villarreal, C. X. & Chan, D. D. Taxonomic Shifts Correlate to Serum Cytokines in an Antibiotics Model to Study the Murine Gut-Joint Axis. 2025.02.15.638396 Preprint at 10.1101/2025.02.15.638396 (2025).

94. Bio-Rad. Bio-Plex Pro Assay 384-Well Protocol Bulletin 7269.

95. Kolde, R. pheatmap: Pretty Heatmaps. (2019).

96. Rohart, F., Gautier, B., Singh, A. & Cao, K.-A. L. mixOmics: An R package for ‘omics feature selection and multiple data integration. PLOS Computational Biology 13, e1005752 (2017).

97. González, I., Déjean, S., Martin, P. G. P. & Baccini, A. CCA: An R Package to Extend Canonical Correlation Analysis. Journal of Statistical Software 23, 1–14 (2008).

