## Supplementary Data 2 for "Metabolites associated with type 2 diabetes and Alzheimer’s disease trigger differential intracellular signaling responses in mouse primary neurons"

### Canonical Correlations

```
[DataSet1] /Users/bbkazu/Desktop/pathway_and_cytokine_cca.sav
```

#### Canonical Correlations Settings

| | Values |
|---|---|
| Set 1 Variables | STAT3 STAT5 ATF2 HSP27 p53 MEK1 MSK1 STAT1 cJun JNK ErkMAPK p38 CREB NFkB Akt p70S6K |
| Set 2 Variables | IL9 MCP1 |
| Centered Dataset | None |
| Scoring Syntax | None |
| Correlations Used for Scoring | 2 |

#### Canonical Correlations

| | Correlation | Eigenvalue | Wilks Statistic | F | Num D.F | Denom D.F. | Sig. |
|---|---|---|---|---|---|---|---|
| 1 | .979 | 23.280 | .008 | 5.914 | 32.000 | 18.000 | <.001 |
| 2 | .904 | 4.460 | .183 | 2.973 | 15.000 | 10.000 | .044 |

H0 for Wilks test is that the correlations in the current and following rows are zero

**Set 1 Standardized Canonical Correlation Coefficients**

| Variable | 1 | 2 |
|---|---|---|
| STAT3 | .102 | .346 |
| STAT5 | .245 | -.424 |
| ATF2 | 1.425 | 1.213 |
| HSP27 | .127 | -1.639 |
| p53 | -.308 | -.338 |
| MEK1 | -.902 | -1.149 |
| MSK1 | -.330 | -1.015 |
| STAT1 | -.259 | -.664 |
| cJun | -.496 | .946 |
| JNK | -1.031 | -2.846 |
| ErkMAPK | .155 | 2.228 |
| p38 | -.020 | .964 |
| CREB | -.816 | -1.072 |
| NFkB | .022 | -.872 |
| Akt | .564 | 2.447 |
| p70S6K | .002 | -.713 |

**Set 2 Standardized Canonical Correlation Coefficients**

| Variable | 1 | 2 |
|---|---|---|
| IL9 | .824 | .695 |
| MCP1 | -.337 | 1.024 |

**Set 1 Unstandardized Canonical Correlation Coefficients**

| Variable | 1 | 2 |
|---|---|---|
| STAT3 | .561 | 1.907 |
| STAT5 | 1.108 | -1.920 |
| ATF2 | 3.358 | 2.857 |
| HSP27 | .097 | -1.253 |
| p53 | -3.444 | -3.786 |
| MEK1 | -1.877 | -2.391 |
| MSK1 | -.662 | -2.035 |
| STAT1 | -.141 | -.362 |
| cJun | -6.105 | 11.647 |
| JNK | -.754 | -2.081 |
| ErkMAPK | .352 | 5.043 |
| p38 | -.084 | 4.045 |
| CREB | -2.212 | -2.907 |
| NFkB | .177 | -7.027 |
| Akt | 1.637 | 7.103 |
| p70S6K | .004 | -1.629 |

**Set 2 Unstandardized Canonical Correlation Coefficients**

| Variable | 1 | 2 |
|---|---|---|
| IL9 | 1.114 | .939 |
| MCP1 | -.505 | 1.535 |

#### Set 1 Canonical Loadings

| Variable | 1 | 2 |
|---|---|---|
| STAT3 | .333 | -.514 |
| STAT5 | .523 | -.285 |
| ATF2 | .112 | .206 |
| HSP27 | -.053 | -.235 |
| p53 | -.114 | -.123 |
| MEK1 | -.513 | .226 |
| MSK1 | .203 | .162 |
| STAT1 | -.188 | .026 |
| cJun | -.696 | .094 |
| JNK | .440 | -.059 |
| ErkMAPK | -.054 | .315 |
| p38 | -.069 | -.027 |
| CREB | .067 | -.220 |
| NFkB | .305 | -.124 |
| Akt | .656 | -.275 |
| p70S6K | -.236 | -.308 |

#### Set 2 Canonical Loadings

| Variable | 1 | 2 |
|---|---|---|
| IL9 | .950 | .312 |
| MCP1 | -.645 | .765 |

#### Set 1 Cross Loadings

| Variable | 1 | 2 |
|---|---|---|
| STAT3 | .326 | -.465 |
| STAT5 | .512 | -.258 |
| ATF2 | .109 | .186 |
| HSP27 | -.052 | -.213 |
| p53 | -.111 | -.111 |
| MEK1 | -.502 | .204 |
| MSK1 | .199 | .147 |
| STAT1 | -.184 | .024 |
| cJun | -.682 | .085 |
| JNK | .431 | -.053 |
| ErkMAPK | -.053 | .284 |
| p38 | -.068 | -.025 |
| CREB | .065 | -.199 |
| NFkB | .299 | -.112 |
| Akt | .642 | -.249 |
| p70S6K | -.231 | -.278 |

#### Set 2 Cross Loadings

| Variable | 1 | 2 |
|---|---|---|
| IL9 | .930 | .282 |
| MCP1 | -.631 | .691 |

#### Proportion of Variance Explained

| Canonical Variable | Set 1 by Self | Set 1 by Set 2 | Set 2 by Self | Set 2 by Set 1 |
|---|---|---|---|---|
| 1 | .126 | .121 | .659 | .632 |
| 2 | .055 | .045 | .341 | .279 |
